# Frequency Chaos Game Representation - Singular Value Decomposition for Alignment-Free Phylogenetic Analysis

**DOI:** 10.1101/2025.04.16.649090

**Authors:** Varsha Achuthan, Deeptangi Mangsuli, Neelam Sinha, Shweta Ramdas

**Affiliations:** Centre For Brain Research, Indian Institute Of Science, India

**Keywords:** Alignment-free Method, Frequency Chaos Game Representation, Singular Value Decomposition, Phylogenetic Tree, Dimension Reduction, Neighbour Joining

## Abstract

This paper introduces ”SVD-FCGR,” a scalable and efficient framework for phylogenetic analysis using Singular Value Decomposition (SVD) on Frequency Chaos Game Representation (FCGR). Unlike traditional MSA techniques, SVD-FCGR handles large datasets with lower computational complexity. It supports both genome-wide and gene-specific analyses, as demonstrated with datasets of Japanese Encephalitis Virus (JEV), Hepatitis B Virus (HBV), Human Immunodeficiency Virus(HIV-1), and Severe Acute Respiratory Syndrome Coronavirus 2 (SARS-CoV-2). For COVID-19, gene-level analysis of surface glycoprotein highlighted mutations affecting viral adaptability, while the envelope gene remained conserved. The method produced detailed phylogenetic trees, surpassing tools like MEGA in resolution and scalability. Validation from the AF Project ranked it 11th among alignment-free methods, confirming its reliability and adaptability.

## 1 INTRODUCTION

Sequence alignment is a fundamental tool in genomic analysis, aiding in the discovery of genetic commonalities, tracing evolutionary relationships, predicting protein structures, and identifying functional domains[1, 2]. Traditional alignment methods are effective for short nucleotide sequences but face challenges with longer sequences[3]. This necessitates the development of alignment-free methods, which are computationally efficient and suitable for large-scale analysis using whole genome sequencing data. [4, 5, 6].

**Figure 1:**
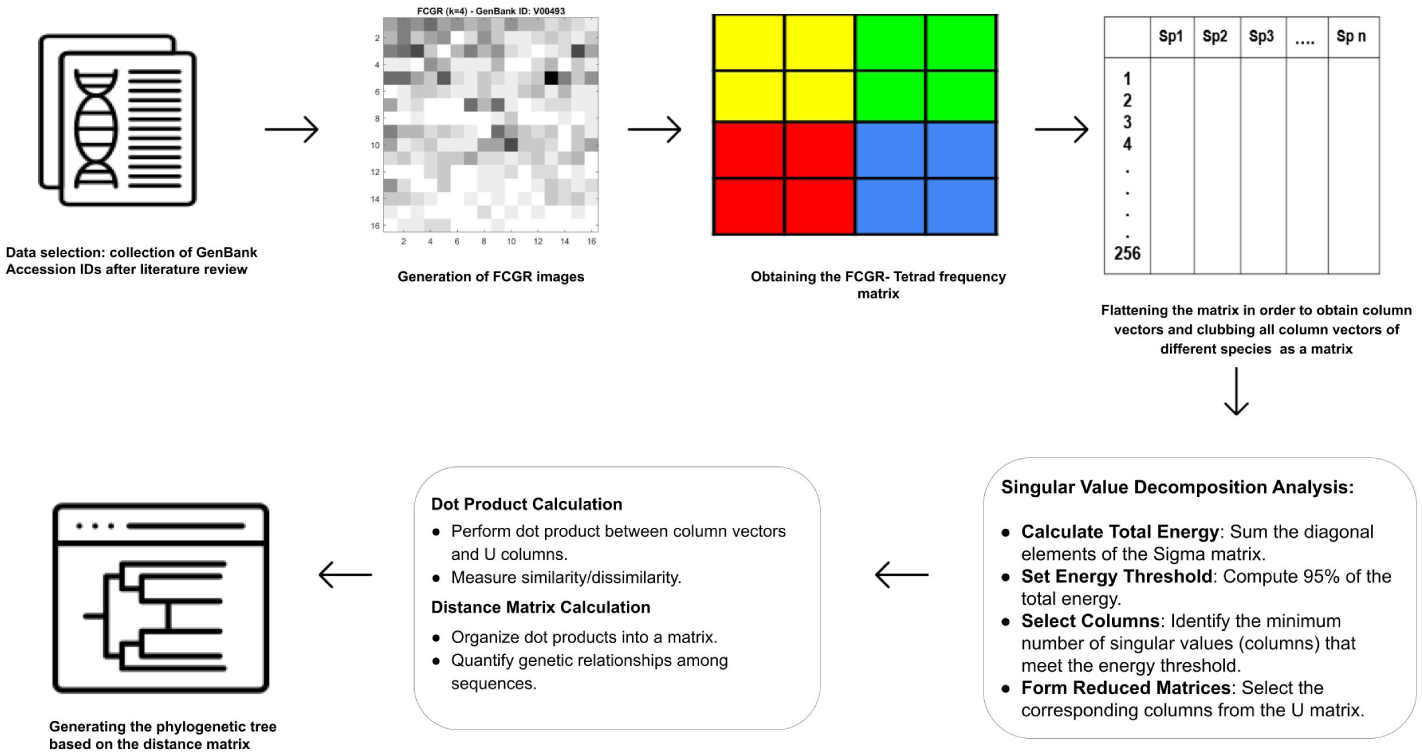
The image illustrates a detailed workflow for generating a phylogenetic tree. The process begins with selecting GenBank Accession IDs after a literature review. This data generates FCGR (Frequency Chaos Game Representation) images. These images are then converted into a Tetrad frequency matrix, flattened to obtain column vectors for different species. Singular Value Decomposition (SVD) analysis is performed on this matrix to calculate total energy, set an energy threshold, select columns, and form reduced matrices. Dot product calculations between column vectors and U columns are used to measure similarity or dissimilarity, and a distance matrix is created to quantify genetic relationships among sequences. Finally, a phylogenetic tree is generated based on the distance matrix.

Alignment-free methods have emerged as powerful tools in computational biology for analyzing genomic sequences without requiring sequence alignment. These methods offer a range of advantages, particularly in terms of computational efficiency and the ability to handle large-scale genomic data[2, 4]. Unlike traditional alignment-based approaches, which rely on explicitly matching nucleotide or amino acid sequences, alignment-free techniques utilize various mathematical and statistical descriptors to capture the essence of sequence composition and structure[1]. Common strategies include k-mer analysis and Frequency chaos game representation (FCGR) [5].

FCGR is employed in fast sequence lookup, constructing phylogenetic trees, and classifying viral genomic data[5]. FCGR reduces ambiguity and offers good granularity, supported by existing research[7]. Each species has a unique Tetrad profile; the identification and quantification of this profile can be applicable in various fields, including meta-genomics, comparative genomics, and genomic signatures[8] to answer questions like species identification, taxonomic classification, primer designing, genome editing,understanding horizontal gene transfer and identifying CpG islands.

## 2 MATERIALS AND METHODS

### 2.1 Frequency Chaos Game Representation (FCGR)

Originally proposed by H. Joel Jeffrey, FCGR is a scale-independent method for genomic sequence comparison[9]. It generates images from nucleotide sequences, capturing fractal properties associated with k-mers (short subsequences of length k).

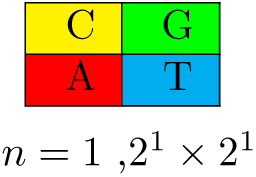

The process involves dividing a DNA sequence into overlapping k-mers, which are then mapped to positions within a 2*^n^ ×* 2*^n^* grid, where *n* represents the k-mer length. The occurrences of each k-mer are counted within their respective grid squares[10]. The resulting image or numeric vector represents the frequencies of all possible k-mers in the original sequence, providing a comprehensive, alignment-free genomic data comparison. The reference figures can be accessed in the supplementary.

### 2.2 Generation of FCGR Images and k-mer matrix construction

The FCGR images, displayed in grayscale, represent the frequency distribution of k-mers within genomic sequences. In this study, a k-mer size of 4 was chosen to create the FCGR images, offering a balanced approach that captures sufficient sequence complexity while maintaining manageable computational demands. Smaller k-mers (e.g., 3-mers) may not capture enough sequence information, leading to oversimplified representations, while larger k-mers (e.g., 5-mers or higher) can introduce excessive computational overhead and result in sparse distributions that are difficult to interpret visually.[10, 11].

Using a k-mer size of 4 strikes an optimal balance in the FCGR images, offering detailed and interpretable visualizations of k-mer frequency distributions within genomic sequences. Since there are four nucleotides—Adenine (A), Guanine (G), Cytosine (C), Thymine (T), or Uracil (U) in RNA—a k-mer size of 4 ensures an unbiased representation, as every possible k-mer can be accurately captured and distinguished within the FCGR framework. [5, 12]. The FCGR image and the Tetrad matrix are fundamentally the same, with the Tetrad matrix acting as the underlying data structure and the FCGR image serving as its visual representation, where grayscale intensity corresponds to k-mer frequency—white for a frequency of 0 and black for the highest frequency. To generate the Tetrad matrices, a sliding window approach is applied to the DNA sequence. Initially, all k-mer counts are set to zero. The process begins by taking four consecutive nucleotides to form a k-mer, which is then assigned to the corresponding cell in the grid matrix. The k-mer frequency count in that cell is incremented by one, and the window shifts forward by one nucleotide to generate the next k-mer. This continues until the entire sequence is traversed, capturing all possible overlapping k-mers.

Once the Tetrad matrices (16 × 16 grids) are generated, they are flattened into 256 × 1 vectors, which are then normalized[13]. Each normalized vector is stored as a column in a larger matrix, with each column representing a different genomic sequence, identified by its species name or GenBank Accession ID. This structured representation facilitates sequence comparison and enables advanced analyses, such as Singular Value Decomposition (SVD), clustering, and phylogenetic studies.

### 2.3 Dimension Reduction

Singular Value Decomposition (SVD) is used for analysis, helping to identify the most significant data components [14]. This step is crucial for simplifying the dataset and making it more manageable for subsequent analysis. First, the total energy is calculated by summing the diagonal elements of the Sigma matrix. An adaptive approach can be taken to set the energy threshold, which can vary depending on species-specific considerations. For example, 95% of the total energy was set as the energy threshold in the proposed SVD-FCGR method, but this percentage can be adjusted as needed to suit different datasets. The choice of using 95% of the total energy as a threshold in Singular Value Decomposition (SVD) is not a one-size-fits-all solution. Different genomic sequences and data sets have unique characteristics, and a fixed energy fraction can either oversimplify or retain too much complexity, potentially affecting the accuracy and relevance of the analysis. Therefore, an adaptive approach to setting the energy threshold is more appropriate. This method allows the threshold to be tailored to the specific features of the dataset, ensuring that the selected number of singular values accurately captures the essential genetic information while minimizing noise and redundancy. By adjusting the energy fraction based on the dataset’s characteristics, the robustness and reliability of phylo-genetic analyses can be enhanced.

The minimum number of singular values that meet this energy threshold is selected, and the corresponding columns from the U matrix are used for further calculations. A dot product is performed between the column vectors representing the different GenBank Accession IDs and the selected columns from the U matrix. These dot products measure the similarity or dissimilarity between the sequences. By organizing these dot products into a matrix, a distance matrix is calculated, quantifying the genetic relationships among the sequences [7].

### 2.4 Data Analysis

Subsequently, hierarchical clustering was performed using the Neighbor Joining (NJ) algorithm. This method constructs a phylogenetic tree based on pairwise distance data, focusing on minimizing the total branch length of the tree. The NJ algorithm iteratively identifies the closest pairs of data points, calculates the shortest branch lengths, and connects them to form clusters. It continues until all data points are merged into a single tree, providing a clear visual representation of the genetic relationships and evolutionary distances.

To cross-verify the results obtained from the SVD-FCGR method, a Multiple Sequence Alignment (MSA) was conducted using MEGA version 12[15]. MSA involves aligning the sequences to identify regions of similarity, which is crucial for understanding evolutionary relationships. The Neighbour Joining algorithm within MEGA was then used to generate another phylogenetic tree based on the aligned sequences. This tree served as a benchmark to validate the results obtained from the SVD-FCGR method.

The phylogenetic trees generated by both methods were then compared to evaluate the accuracy and reliability of the SVD-FCGR method in capturing phylogenetic relationships. This comparative analysis reinforced the confidence in the new method, demonstrating its potential for broader applications in genomic analysis

### 2.5 Flexible Phylogenetic Analysis for Genomes and Genes

This approach is versatile, allowing for analysis of both whole genome sequences and specific CDS (Coding DNA Sequences). For example, we analyzed four viruses using this method. For HIV-1-1, JEV, and HBV, we utilized their whole genomes to generate phylogenetic trees, ensuring comprehensive representation of their genetic diversity. In the case of COVID-19, we performed gene-level analysis for two specific genes, demonstrating the flexibility of this workflow. Users are prompted to choose between comparing the entire genome or selecting a particular gene from a displayed gene table, allowing for tailored and precise comparisons to support diverse research objectives. This adaptability enhances the utility of the method across various viral datasets.

## 3 DATA SELECTION

### 3.1 Japanese Encephalitis Virus

Japanese encephalitis (JE) is a severe neurological disease caused by the Japanese encephalitis virus (JEV), primarily transmitted by *Culex* mosquitoes, with birds as intermediate hosts and pigs as amplifying hosts [16, 17, 18]. JE has a high fatality rate, and survivors may suffer lasting neurological or psychological damage, including aphasia, paralysis, and consciousness impairment [19].

JEV is an enveloped, neurotropic virus with a single-stranded RNA genome (11,000 base pairs), containing three structural proteins—core protein (C), membrane protein (M), and glycosylated envelope protein (E)—and seven non-structural proteins: NS1, NS2A, NS2B, NS3, NS4A, NS4B, and NS5 [20, 21]. It crosses the blood-brain barrier, replicates in neurons, and causes neurolysis, leading to cerebral edema, brain-stem herniation, and death.

The E protein induces virus-neutralizing antibodies, providing protective immunity [20]. JEV strains are classified into five genotypes (GI-V) based on E gene phylogenetic analysis [22]. Most Indian isolates belong to GIII, while GI has recently been reported in Gorakhpur, Uttar Pradesh [16, 19]. Genotypes are distributed as follows: GI (Southeast Asia, Australia, India), GII (Southern Thailand, Malaysia, Indonesia, Australia), GIII (Southeast Asia, Japan, India), GIV (Indonesia), and GV (Singapore) [23, 20, 24].

#### 3.1.1 Dataset Selection for JEV

A total of 3,541 Japanese Encephalitis Virus (JEV) sequences were obtained from the NCBI Virus database [25]. This dataset included 384 complete sequences and 3,157 partial sequences. From the complete sequences, 18 isolated from Indian species were selected, forming a small dataset for analysis, ensuring the quality and specificity of the genetic data [26]. The GenBank Accession IDs of these sequences are provided in the Supplementary Information.

### 3.2 Human Immunodeficiency Virus

Human Immunodeficiency Virus (HIV-1) is a retrovirus with a compact genome of approximately 9.75 kilobases, comprising two single-stranded RNA molecules linked in a dimeric structure [[27]]. The virus is enveloped in a lipid bilayer embedded with glycoproteins gp120 and gp41, which facilitate attachment and fusion to CD4+ T cells, the primary targets of HIV-1 [[28]]. Inside the conical capsid, essential enzymes such as reverse transcriptase, integrase, and protease work together to reverse transcribe the viral RNA into DNA and integrate it into the host genome, enabling persistent infection and replication [[29]].

HIV-1 is marked by remarkable genetic diversity driven by high mutation rates and recombination during replication, which pose challenges for vaccine development and therapeutic intervention [[27]]. Its structure and replication mechanisms have made it one of the most studied viruses in virology, providing insights into immunology and structural biology. These discoveries have informed strategies to mitigate the global impact of HIV-1, which continues to affect approximately 39 million people worldwide [[29]].

#### 3.2.1 Dataset Selection for HIV-1

A total of 16 HIV-1 subtype C strains were selected from the LANL HIV-1 Sequence Database [[30]]. The dataset comprised 8 strains from India and 8 strains from South Africa, ensuring diverse geographic representation. The selection focused on subtype C due to its prevalence and significance in global HIV-1-1 research.

### 3.3 Hepatitis B Virus

Hepatitis B Virus (HBV) is about 3,200 base pairs long, making it one of the smallest DNA viruses [[31]]. Its compact genome encodes multiple essential proteins, including the surface antigen (HBsAg), core antigen (HBcAg), polymerase, and X protein, which are essential for replication and the virus’s ability to persist in the host [[32]]. HBV belongs to the Hepadnaviridae family, utilizing a unique replication strategy involving both DNA and RNA intermediates. This replication process, driven by reverse transcriptase activity, contributes to genetic variability and resistance to antiviral therapies [[32]].

Globally, HBV infects over 250 million people and causes approximately 900,000 deaths annually, primarily due to complications such as cirrhosis and hepatocellular carcinoma [[33]]. Transmission occurs via blood, sexual contact, or perinatally during childbirth. Despite the availability of effective vaccines, HBV remains a public health concern, particularly in low- and middle-income countries where access to healthcare is limited [[33]]. Recent research advancements focus on RNA interference and immune modulators, offering hope for functional cures by targeting sustained viral suppression and immune control [[32]].

#### 3.3.1 Dataset Selection for HBV

A total of 78 Hepatitis B Virus (HBV) sequences were randomly selected from the HBVdb database [[34]]. This dataset included four subtypes: A, B, D, and G, with 20 sequences each for subtypes A, B, and D, and 18 sequences for subtype G. The random selection ensured diverse representation of HBV genetic variants, encompassing major subtypes for comprehensive analysis. Detailed GenBank accession IDs for these sequences are provided in the supplementary information

### 3.4 SARS-CoV-2

SARS-CoV-2, the virus behind the COVID-19 pandemic, is a positive-sense single-stranded RNA virus first identified in Wuhan, China, in 2019. It spreads primarily via respiratory droplets and enters cells by binding to the angiotensin-converting enzyme 2 (ACE2) receptor [35, 36]. Symptoms range from mild to severe, with older adults and those with comorbidities at higher risk. Vaccination remains a key tool in reducing severe cases and fatalities [37, 38].

The B.1.617.1 variant (Kappa), first reported in India, carries mutations (L452R, E484Q, P681R) that enhance cell entry, immune evasion, and transmissibility [37]. While less transmissible than the Delta variant, it has rapidly spread in regions where it emerged [38]. Vaccines largely remain effective, offering significant protection against severe disease, though with slight reductions in efficacy [37]. Containment measures include enhanced surveillance, genomic sequencing, and public health initiatives, emphasizing the need for strong infrastructure to manage SARS-CoV-2 variants [19, 39].

#### 3.4.1 Dataset Selection of SARS-CoV-2

A dataset comprising of 30 strains, representing Indian variants of the B.1.617.1 lineage, was selected from the NCBI Virus database [25]. The samples were randomly chosen and downloaded using NCBI filters for sequence completeness, quality scores, and host species. To ensure reliable and accurate analysis, only complete sequences with high quality scores were included. The GenBank Accession IDs for these strains are provided in a table in the supplementary information.

## 4 Results

### 4.1 Generation of FCGR Images

The algorithms for generating the Frequency Chaos Game Representation (FCGR) images and the Tetrad matrix were developed. The FCGR images, displayed in gray scale, represent the frequency distribution of k-mers within the sequences. The results presented below demonstrate the efficacy of the proposed SVD-FCGR Method sequences.[5, 14]

**Figure 2:**
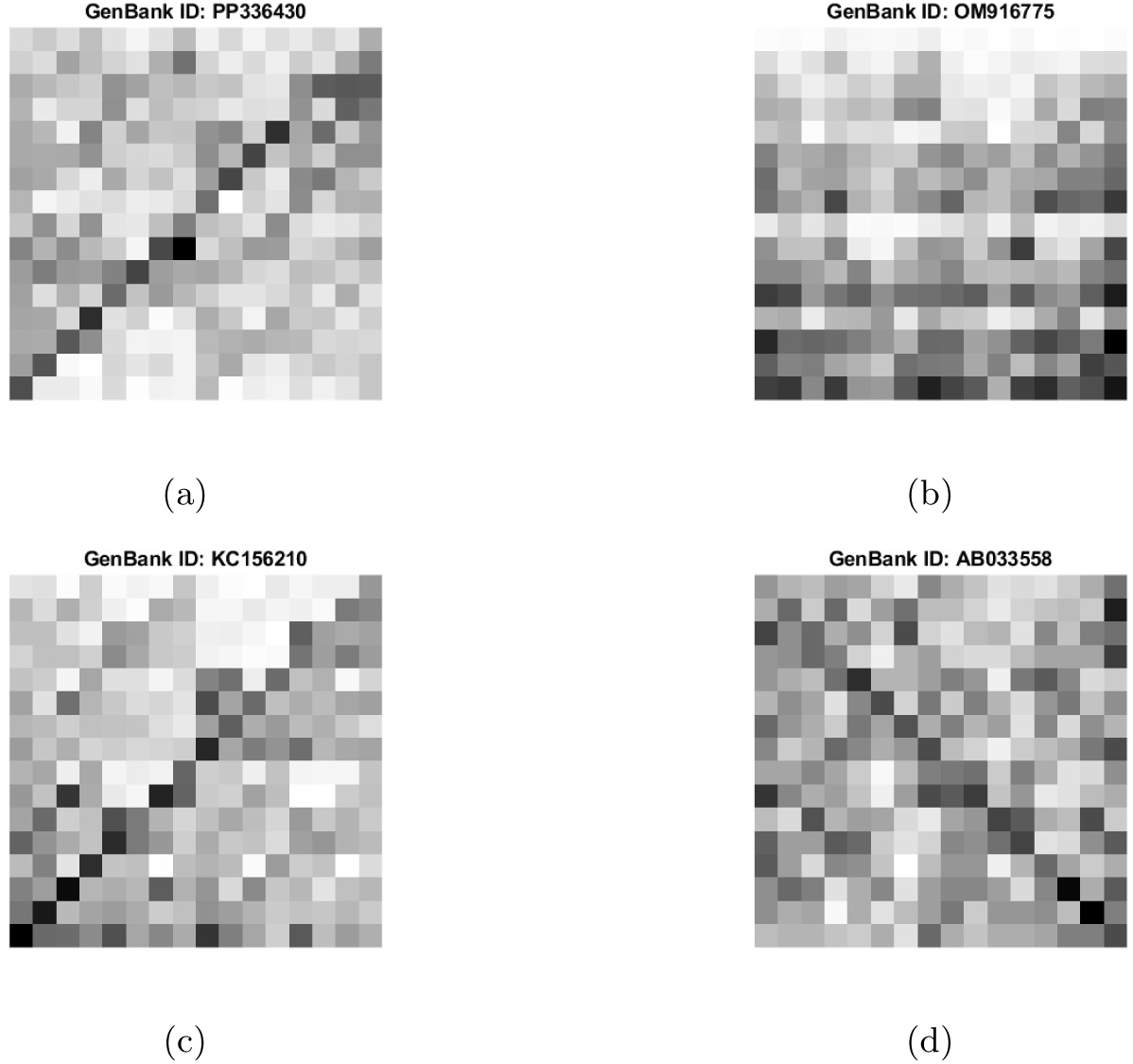
The images represent the Frequency Chaos Game Representation (FCGR) of 4 selected samples from the JEV dataset, each displayed in grayscale. Each FCGR image captures the frequency distribution of k-mers within the genomic sequences, highlighting variations and patterns unique to each sample. The GenBank IDs of the samples considered are labeled as follows: (a)PP336430, (b)OM916775, (c)KC156210, (d)AB033558.

The phylogenetic tree obtained from the proposed SVD-FCGR method and the one obtained from the sequence alignment tool are compared.

**Figure 3:**
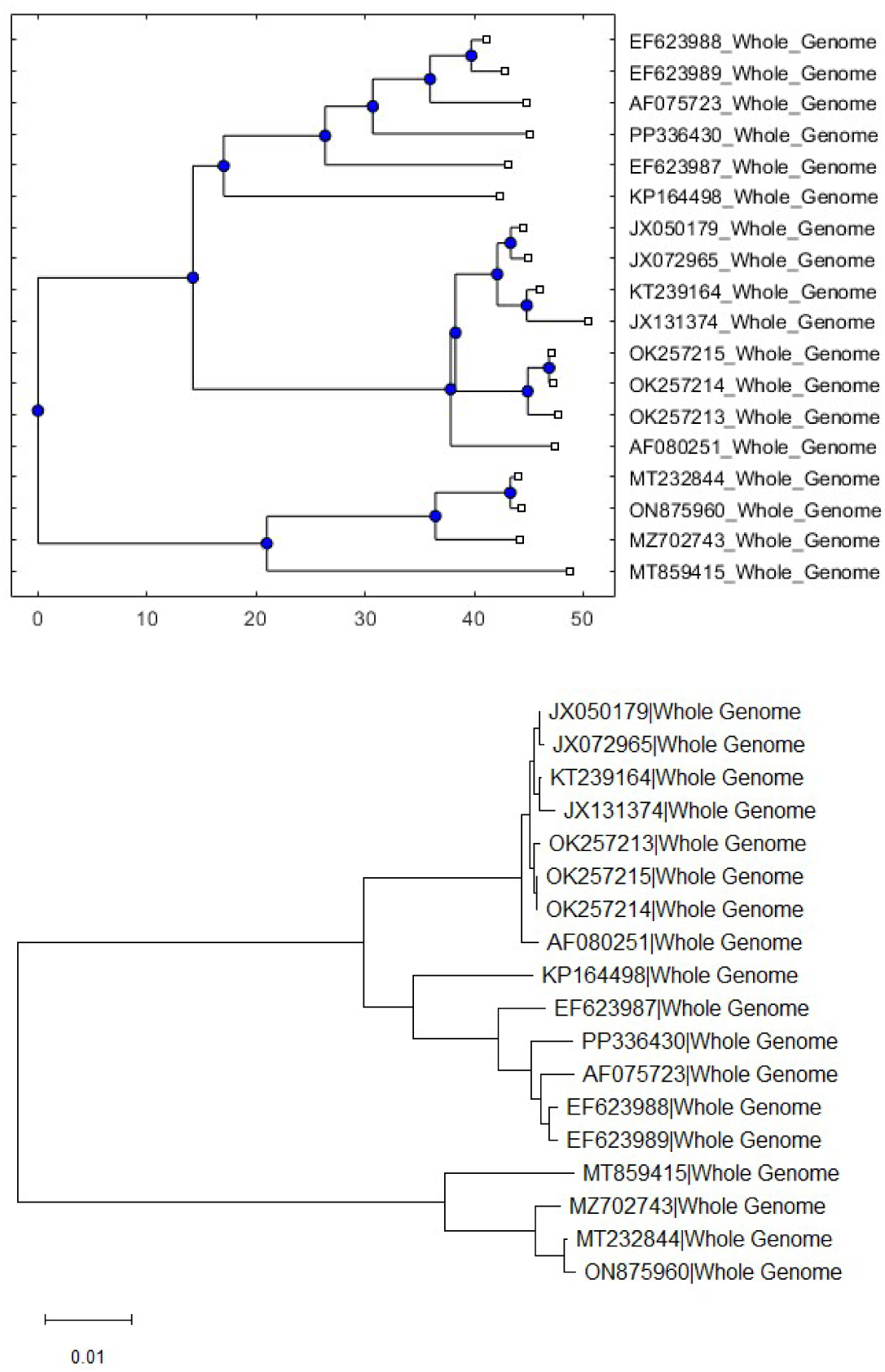
The image above shows the phylogenetic tree obtained from the proposed SVD-FCGR method for Japanese Encephalitis virus dataset. The image below depicts the phylogenetic tree generated using MEGA software for the same dataset

The identical phylogenetic trees obtained using the SVD-FCGR method and MEGA 12 software show consistent branching patterns, indicating that the evolutionary relationships among sequences were accurately preserved by both methods. The topology of the trees, including the arrangement and divergence of branches, confirms that the two approaches align in reconstructing genetic relationships within the dataset.

**Figure 4:**
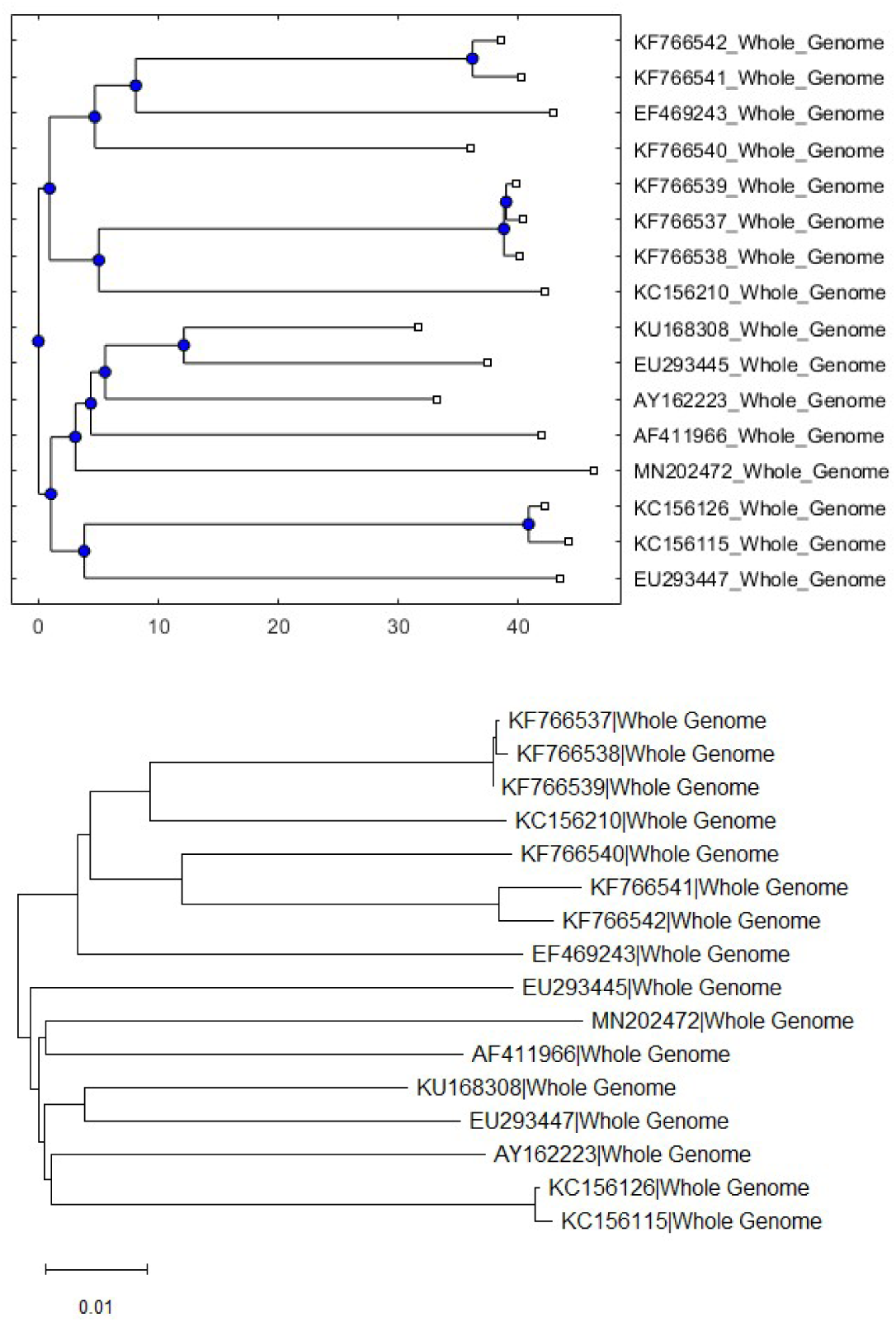
The image above shows the phylogenetic tree obtained from the proposed SVD-FCGR method for HIV-1 dataset. The image below depicts the phylogenetic tree generated using MEGA software for the same dataset

The phylogenetic tree generated for the 16 HIV-1 subtype C strains demonstrates distinct branching patterns, clearly separating the sequences based on their geographic origins. The 8 strains from India and the 8 strains from South Africa form two separate clusters, despite belonging to the same subtype. This segregation highlights subtle genetic variations between the regions, providing insight into the evolutionary divergence of subtype C strains from different geographic locations. The ability of the method to distinguish these patterns underscores its effectiveness in accurately representing genetic relationships across diverse datasets.

**Figure 5:**
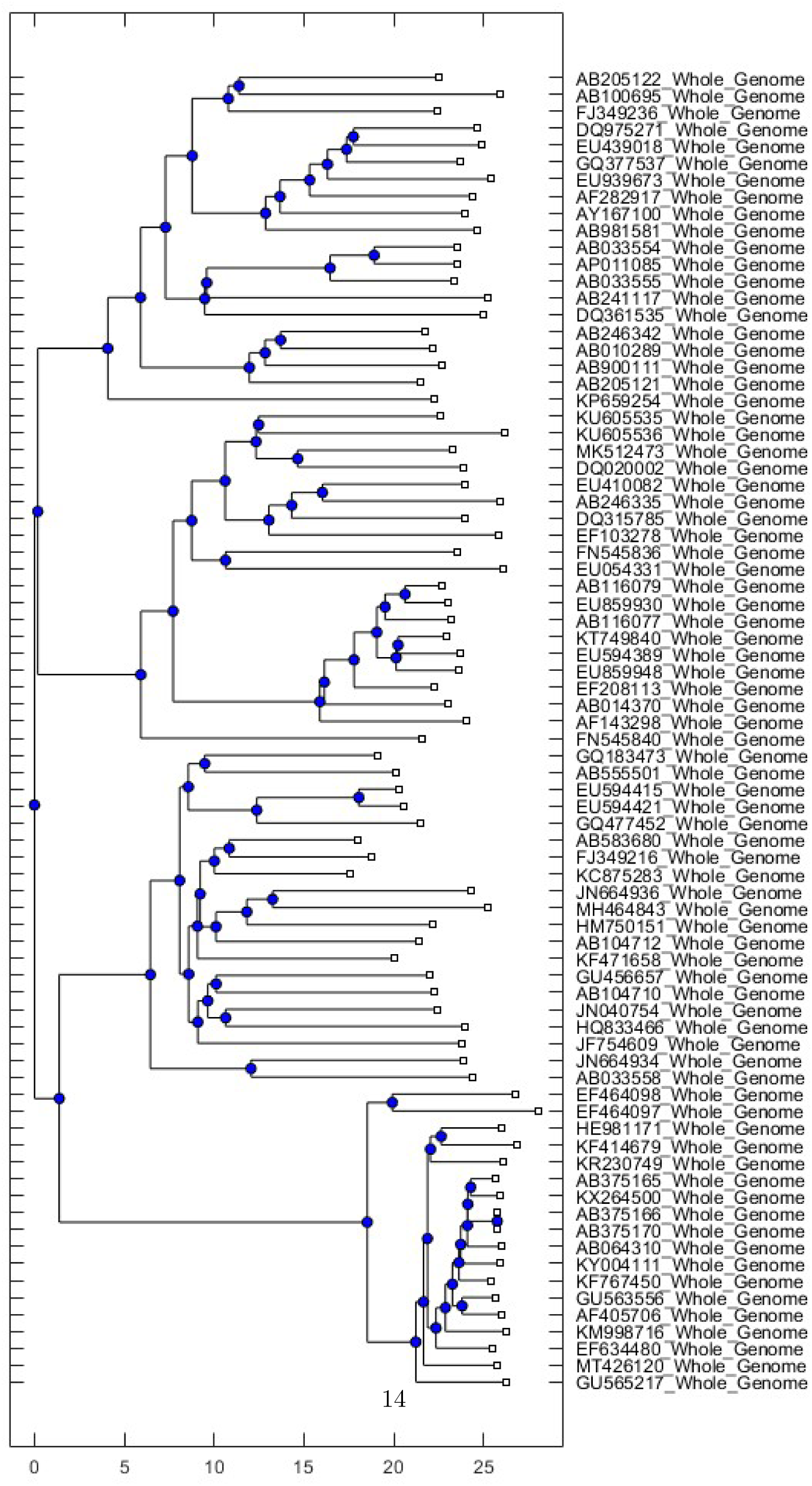
The phylogenetic tree obtained from the proposed SVD-FCGR method for the HBV dataset.

**Figure 6:**
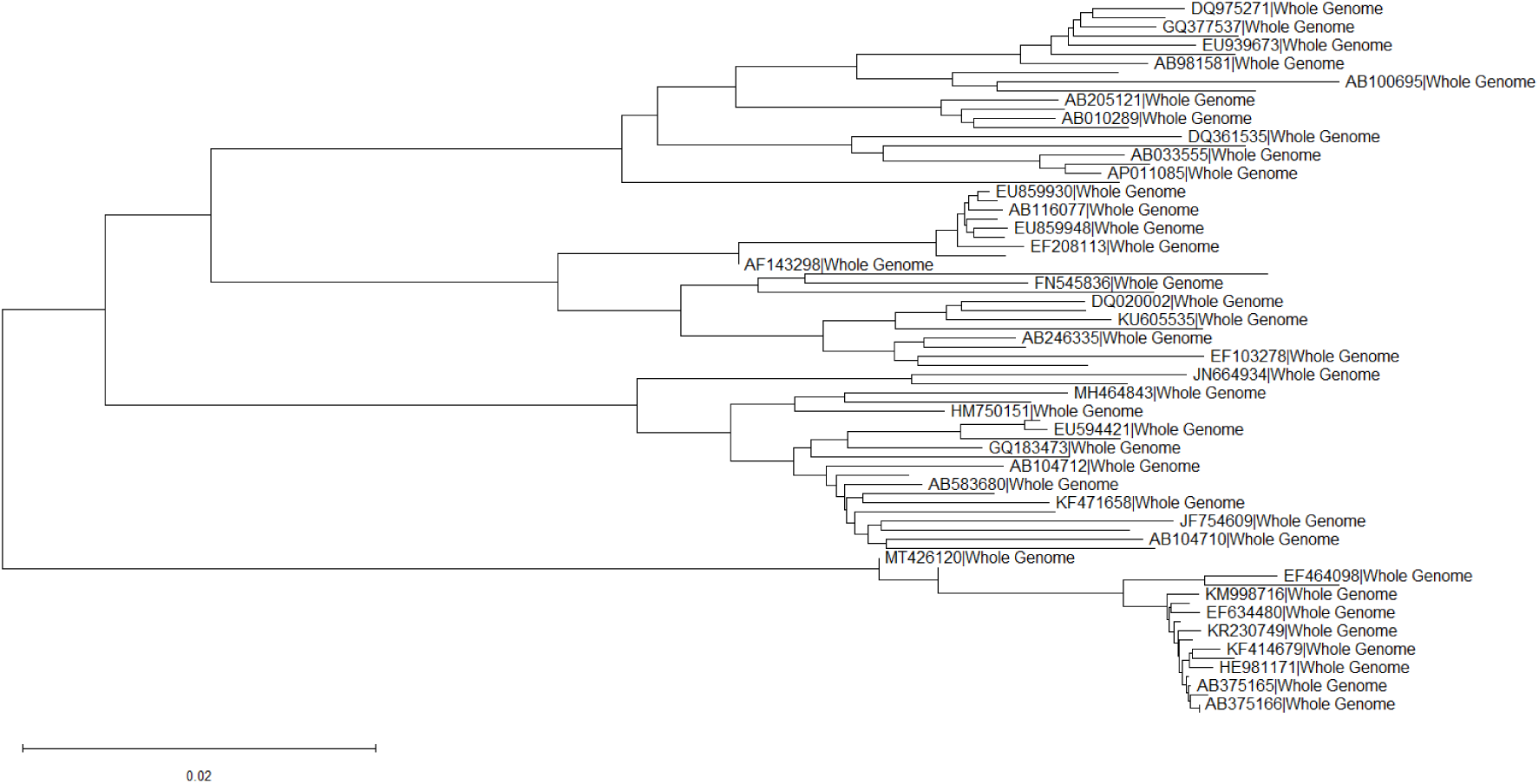
The phylogenetic tree generated using MEGA software for the HBV dataset.

The phylogenetic tree generated using our method successfully classified the substrains—A, B, D, and G—into distinct clusters, allowing verification of their genetic relationships. With our approach, we were able to take as many samples as needed, resulting in 78 unique branches that fully captured the diversity within the dataset. In contrast, the tree generated using MEGA software displayed only 39 branches, failing to account for a significant portion of the data. This demonstrates the scalability of our method, capable of handling large datasets and accurately representing complex evolutionary patterns, making it highly suitable for comprehensive analyses.

**Figure 7:**
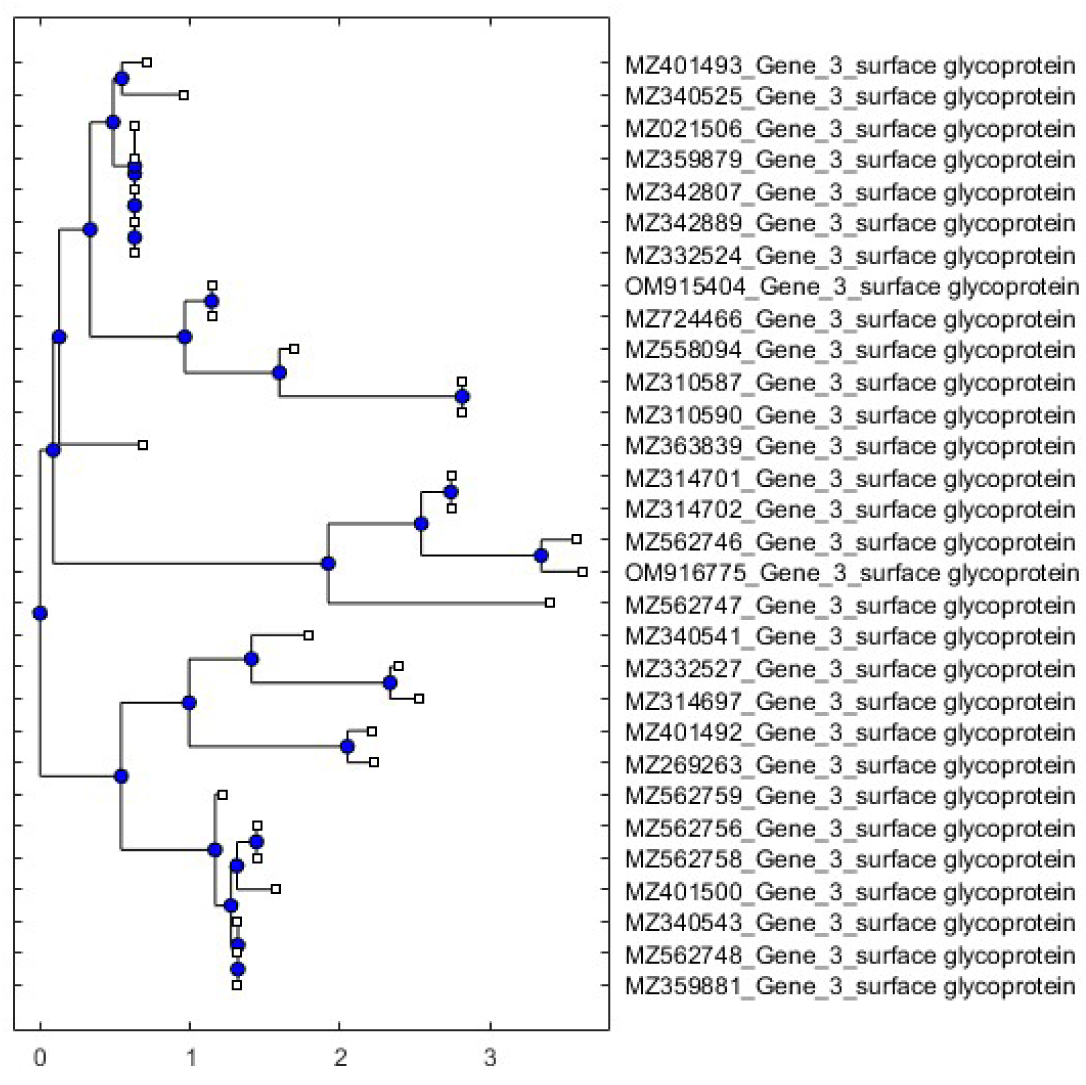
The phylogenetic tree obtained from the proposed SVD-FCGR method for the surface glycoprotein gene of the COVID-19 dataset.

The surface glycoprotein plays a pivotal role in the adaptability and evolution of the COVID-19 virus. Mutations in this protein can significantly alter its structure, impacting the ability of the virus to bind to host cell receptors, such as ACE2, and facilitate entry into cells. Such changes may give rise to new viral strains with enhanced transmissibility or altered immune escape mechanisms, leading to the emergence of variants capable of evading neutralizing antibodies or adapting to diverse host environmentsThe phylogenetic tree generated from the analysis of the surface glycoprotein gene (3822 bp) highlights these evolutionary changes, providing insights into how genetic variations in this region may affect the virus’s virulence and transmission dynamics. This adaptability in gene-level analysis demonstrates the utility of our method in uncovering crucial patterns in viral evolution.

**Figure 8:**
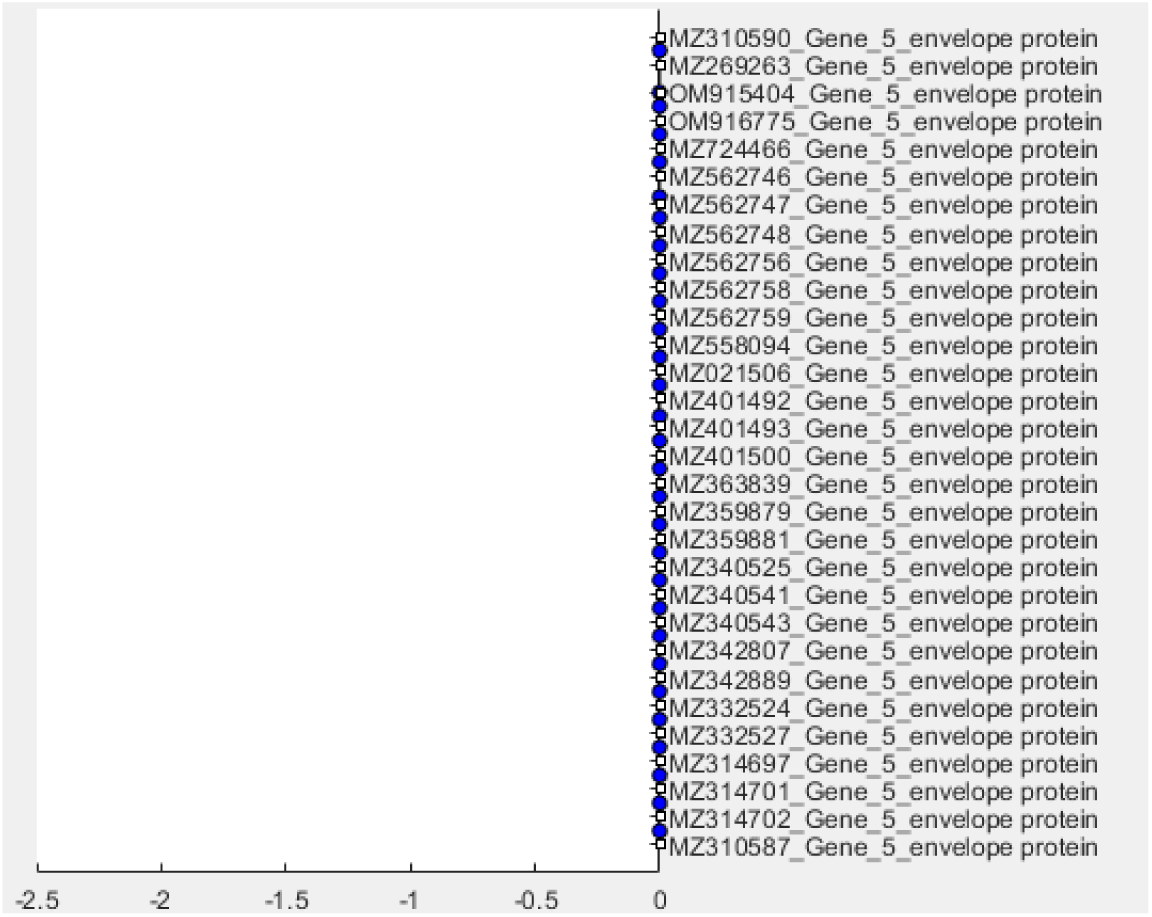
The phylogenetic tree obtained from the proposed SVD-FCGR method for the envelope gene of the COVID-19 dataset.

In contrast, the phylogenetic tree for the envelope gene, which has a length of 228 base pairs, showed no variation at all. All sequences were perfectly identical, resulting in a tree where no differentiation could be observed. To confirm this, a sequence alignment was performed, yielding a 100% match across all the sequences. This confirms that there are no mutations in this region of the CDS, indicating that the envelope gene remains conserved across the dataset. This lack of variability suggests its critical functional role, which may not tolerate mutations, ensuring the structural or functional integrity of the virus. minor disagreements between the trees can occur due to inherent differences in the methodologies. Alignment-based methods, which rely on aligning nucleotide or amino acid sequences, consider specific genetic changes such as substitutions, insertions, and deletions [15]. This process provides detailed insights into sequence evolution but can be computationally intensive and prone to errors with highly divergent sequences. On the other hand, alignment-free methods, which use mathematical and statistical descriptors like k-mer analysis or Frequency Chaos Game Representation (FCGR), capture the overall composition and structure of sequences without requiring explicit alignment [37].

**Table 1:** Elapsed time for running the code on different datasets, demonstrating the scalability of the approach.

| Dataset | Number of Samples | Base Pair Range | Elapsed Time in SVD-FCGR (seconds) | Elapsed Time in MEGA12 (seconds) |
| --- | --- | --- | --- | --- |
| JEV samples | 18 | 10,800 to 11,000 | 36.091412 | 841.05 |
| HIV-1 samples | 16 | 9,600 to 9,800 | 25.647679 | 519.41 |
| HPV samples | 78 | 3,100 to 3,300 | 99.974631 | 1140.86 |
| SARS-CoV-2 samples surface glycoprotein | 30 | 3,822 | 45.03680 | 235.241 |
| SARS-CoV-2 samples envelope protein | 30 | 228 | 33.7880 | 40.659 |

While these methods are computationally efficient and can handle large-scale genomic data, they may overlook some nuances of sequence evolution that alignment-based methods capture. Consequently, while both approaches aim to reconstruct evolutionary relationships, their different strategies and underlying assumptions can lead to slight variations in tree topology [5]. Shared branches indicate common ancestry or relatedness among the entities represented in both trees. While both methods yield similar trees, they offer complementary insights. Pairwise distances emphasize overall similarity/dissimilarity, whereas sequence alignment considers specific genetic changes such as substitutions, insertions, and deletions [37, 38].

### 4.2 Validation

#### Sensitivity Test

To assess the proposed SVD-FCGR method’s sensitivity, the k-mer count was adjusted to k=8. Subsequently, a 256x256 FCGR Tetrad matrix was obtained, and the same procedure was followed for 18 JEV sequences. It was observed that the choice of k significantly impacts the size of the FCGR matrix. For k=4, a smaller 16x16 matrix was obtained, while for k=8, the matrix expanded to 256x256[5]. However, due to computational constraints, for k=8, an economy SVD was used, resulting in a larger U matrix (65536x18)[14]. In addition, the Percentage Identity Matrix (PID) along with the distance matrix were compared to understand the trend between the similarity and dissimilarity. These matrices provide insights into the similarity and dissimilarity among the data points[7]. To standardize the values, z-score normalization was applied, ensuring that the mean is 0 and the standard deviation is 1.

**Figure 9:**
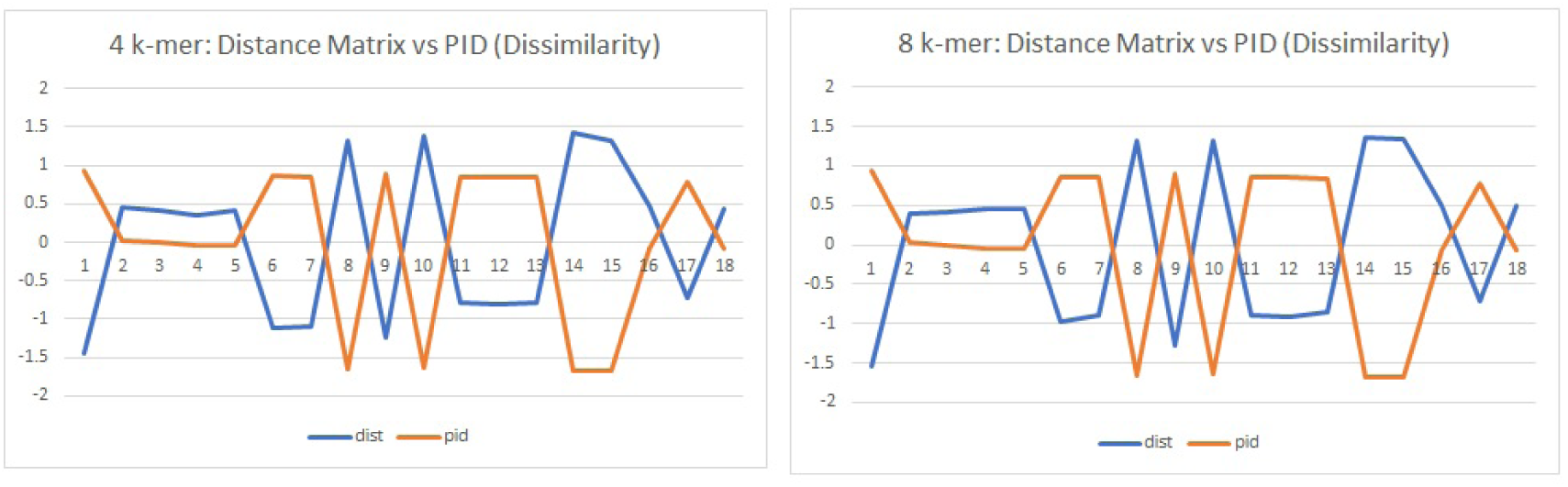
Dissimilarity Trend: The values obtained through k-mer=4 and k-mer=8 distance matrix and Percentage Identity matrix were plotted against each other to show the dissimilarity observed, emphasizing the fact that the proposed SVD-FCGR method is independent of the k-mer length.

**Figure 10:**
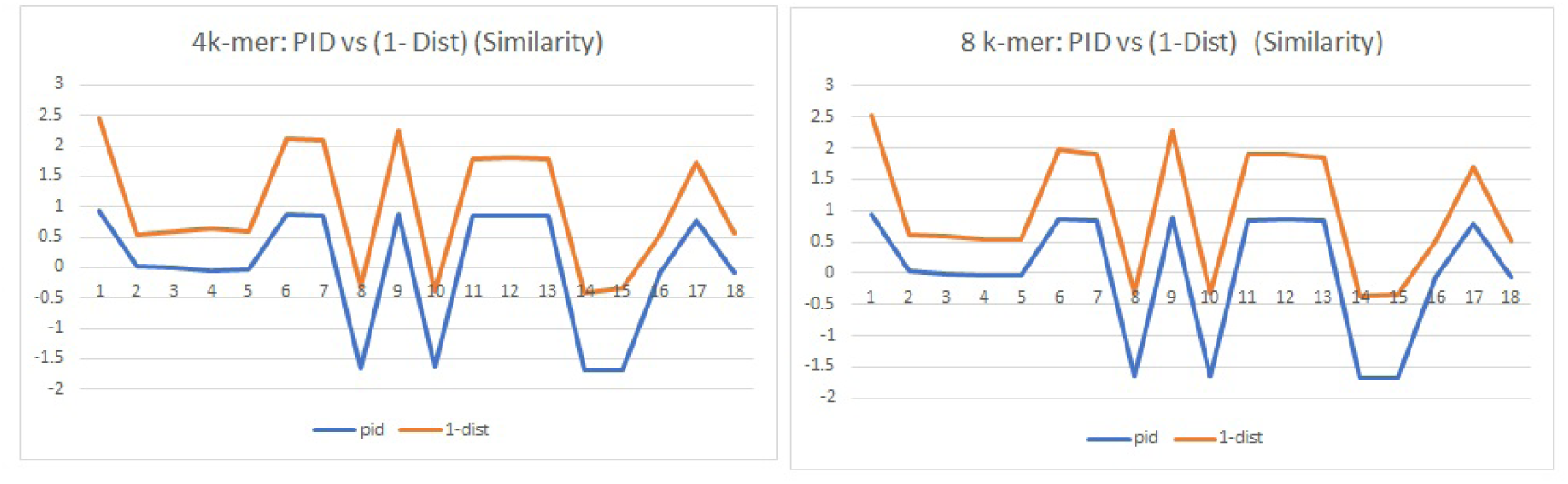
Similarity Trend: The values obtained through k-mer=4 and k-mer=8 Percentage Identity matrix and (1-distance matrix) were plotted against each other to show the similarity observed emphasizing the fact that the proposed SVD-FCGR method is independent of the k-mer length

While analyzing the plots, it is notable that the trend remains consistent even when the k-mer count changes from k=4 to k=8 for both similarity and dissimilarity analyses. This consistency suggests that the method is robust and that, despite altering the k-mer length, the underlying patterns persist, reinforcing the reliability of our approach. This provides a promising alternative to traditional sequence alignment methods

### 4.3 Discussion

In contrast, the SVD-FCGR method successfully generated phylogenetic trees regardless of dataset size. Our algorithm efficiently managed extensive data, overcoming the limitations of traditional software. It was observed that significant differences existed between the software’s output and SVD-FCGR method’s phylogenetic tree. Our approach demonstrated superior efficiency, making it suitable for large-scale analyses. It’s important to note that a phylogenetic tree obtained through an alignment-free method, doesn’t need to match exactly with one obtained through multiple sequence alignment (MSA).

Both methods have strengths and limitations and can produce different results depending on the data and specific algorithms used. The differences in the underlying assumptions and algorithms can lead to variations in the resulting phylogenetic trees. However, both methods aim to reconstruct the evolutionary history of the sequences; the high degree of consistency between the two trees, especially in well-supported clades, reflects the ability of both classes of methods to capture this underlying history.

## 5 Validating the method using AF Project

### Phylogenetic Tree Comparison Metrics

To evaluate the accuracy of our phylogenetic tree analysis using the SVD-FCGR method, we utilized the AF Project website [40]. The website provided several key metrics to compare our method with other alignment-free approaches, namely the Robinson-Foulds (RF) distance, Normalized Robinson-Foulds (nRF) distance, and Normalized Quartet Distance (nQD)

The fish mitochondrial dataset on the AF Project test platform pertains to 25 assembled genomes of fish species from the suborder Labroidei. This dataset was taken from Fisher et al., 2013. The dataset includes 25 FASTA files, each representing a different mitochondrial genome of these fish species. The genomes were sequenced to provide insights into the evolutionary relationships and genetic diversity within this group of fish [41]. The benchmark procedure on the AF Project platform evaluates the accuracy of alignment-free distance measures in reconstructing species phylogeny based on whole genome sequences. Users can download the dataset, run their methods on the provided sequences, and submit their predictions for performance evaluation [40]

The nRF distance, normalized between 0 and 1, allows for easier comparison between methods. Our method’s nRF value was 0.50, suggesting a moderate level of similarity to the benchmark phylogenetic tree [3].

The nQD metric, which assesses the differences in quartet splits, provided an nQD value of 0.1873 for our method. This relatively low value indicates good alignment with the benchmark relationships.

Our SVD-FCGR method was ranked 11th overall on the AF Project website, demonstrating its effectiveness and reliability in reconstructing phylogenetic relationships.

Below is the phylogenetic tree generated using the alignment-free method from the AF Project website. The tree visually represents the genetic relationships among the fish species in our dataset.

**Figure 11:**
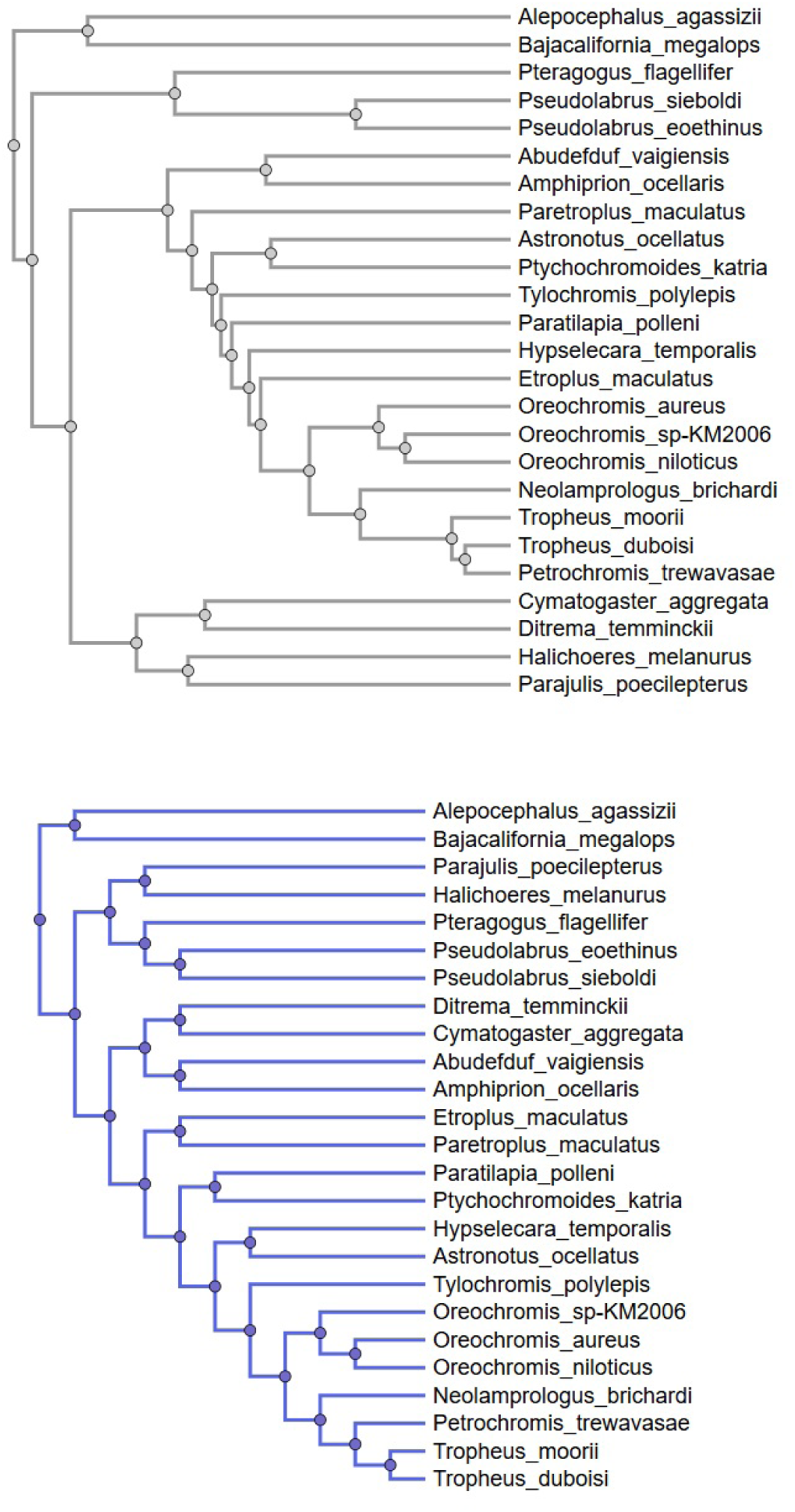
Phylogenetic trees generated using the alignment-free method from the AF Project website. The tree on the top represents the results from our SVD-FCGR method, while the tree on the bottom is the reference tree. This comparison highlights the genetic relationships among the fish species in our dataset. Comparison metrics (nRF, nQD) demonstrate the accuracy and reliability of our method.

## 6 Limitations

The early stages of developing this method show promising results, but there is significant room for refinement and enhancement. Future efforts should focus on improving the algorithm to enhance tree-building capabilities and extend the method’s applicability to larger and more complex datasets. Furthermore, although the proposed SVD-FCGR method demonstrates potential in analyzing phylogenetic relationships with a limited set of genomic sequences, the relatively small dataset used in this study highlights the need for a more comprehensive evaluation. Expanding the dataset in future research could provide a more robust and reliable assessment of the method’s efficacy.

Despite these limitations, the initial results demonstrate the potential of the proposed SVD-FCGR method, offering a promising alternative to traditional alignment-based techniques for large-scale phylogenetic analysis.

## 7 Conclusion

Singular Value Decomposition (SVD). The SVD-FCGR method demonstrated significant advantages over traditional Multiple Sequence Alignment (MSA) techniques, particularly in handling extensive datasets with efficiency and accuracy. The ability to construct phylogenetic trees from extensive datasets has far-reaching implications for research. Traditional MSA often struggles with large datasets, leading to increased computational time and potential inaccuracies.

In contrast, our approach efficiently handles extensive data without compromising on accuracy. This robustness makes the SVD-FCGR method more suitable for large-scale phylogenetic analyses, ensuring reliable tree construction even with complex datasets. The success of the SVD-FCGR method highlights its potential applications and motivates further exploration in this area. Our findings underscore the robustness and reliability of this SVD-FCGR method, making it a viable alternative for large-scale phylogenetic analyses. The ability to construct phylogenetic trees from extensive datasets without compromising accuracy facilitates deeper insights into evolutionary relationships.

## 8 Future work

Expanding dataset diversity by incorporating a wider range of genomic sequences from different species and geographical locations can further validate the method. Additionally, enhancing algorithm efficiency is crucial for handling larger datasets without compromising accuracy. Performing extensive comparative studies with other alignment-free methods will help evaluate relative performance and identify potential areas for improvement.

## 9 Abbreviations

FCGR: Frequency Chaos Game Representation
JEV: Japanese Encephalitis Virus
MSA: Multiple Sequence Alignment
SVD: Singular Value Decomposition
PID: Percentage Identity Matrix
NJ: Neighbour Joining
HIV-1: Human Immunodeficiency Virus
HBV: Hepatitis B Virus
SARS-CoV-2: Severe Acute Respiratory Syndrome Coronavirus 2.

## 10 Declarations

### 10.1 Ethics approval and consent to participate

We confirm that we have read the Journal’s position on issues involved in ethical publication and affirm that this report is consistent with those guidelines.

### 10.2 Consent for publication

We hereby consent to the publication of the manuscript, ”Frequency Chaos Game Representation - Singular Value Decomposition for Alignment-Free Phylogenetic Analysis” in BMC Bioinformatics, is subject to its acceptance.

### 10.3 Availability of data and materials

All the datasets have been added in the supplementary information section. This includes NCBI accession numbers of the samples that are considered in the dataset used for analysis. The raw data supporting the conclusions of this article will be made available by the authors, without undue reservation, to any qualified researcher.

### 10.6 Authors’ contributions

V.A. and D.M. contributed significantly to the conceptualization of the study and the development of its methodology. They were instrumental in the software development, performing formal analyses,visualisation, investigation, and managing data curation. Additionally, V.A. and D.M. were responsible for writing the original draft and took an active role in reviewing and editing the manuscript to ensure its quality and coherence.N.S. played a pivotal role in the conceptualization and validation of the research, providing valuable resources and overseeing the project administration. N.S.’s supervision ensured the project adhered to its objectives and timelines. N.S. also contributed to the manuscript’s review and editing, enhancing its overall clarity and impact.S.R. was involved in the validation of the research, bringing critical insights to ensure the accuracy and reliability of the findings. S.R. also assisted in the manuscript’s review and editing, contributing to its final refinement.All authors collectively ensured the development and completion of the research project, and they reviewed and approved the final manuscript, reflecting their substantial contributions and collaboration.

## Supplementary Information

**Figure S1:**
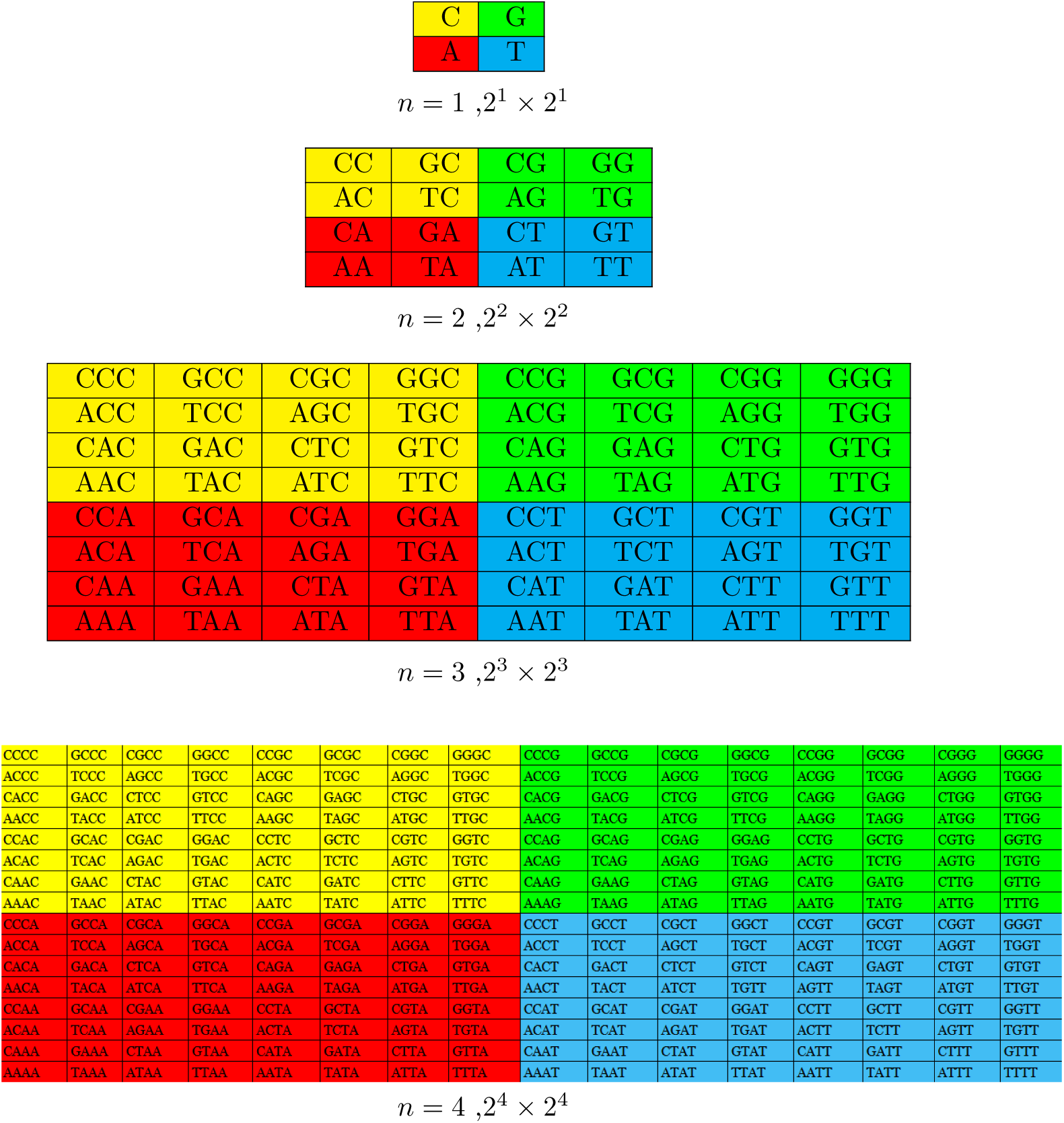
Building the FCGR - Tetrad Frequency Matrix: Each k-mer’s frequency is organized into a matrix, with rows and columns representing specific k-mers. Matrices for k-mers of sizes 1, 2, 3, and 4 are constructed. The matrix cells show how often each k-mer appears in the sequence. Each k-mer is mapped to a cell, and its occurrences are counted. If a k-mer appears multiple times, the corresponding cell’s value increases accordingly.

**Table 2:** List of the 18 Japanese Encephalitis Virus (JEV) samples chosen from the NCBI database.

| Sl no | GenBank Accession ID |
| --- | --- |
| 1 | AF075723 |
| 2 | AF080251 |
| 3 | EF623987 |
| 4 | EF623988 |
| 5 | EF623989 |
| 6 | JX050179 |
| 7 | JX072965 |
| 8 | MT232844 |
| 9 | KT239164 |
| 10 | MZ702743 |
| 11 | OK257215 |
| 12 | OK257214 |
| 13 | OK257213 |
| 14 | MT859415 |
| 15 | ON875960 |
| 16 | PP336430 |
| 17 | KP164498 |
| 18 | JX131374 |

**Table 3:** List of the 16 Human Immunodeficiency Virus samples chosen from the LANL HIV-1 Sequence Database.

| Sl no | GenBank Accession ID |
| --- | --- |
| 1 | EF469243 |
| 2 | KC156210 |
| 3 | KF766540 |
| 4 | KF766541 |
| 5 | KF766542 |
| 6 | KF766539 |
| 7 | KF766537 |
| 8 | KF766538 |
| 9 | KU168308 |
| 10 | MN202472 |
| 11 | KC156126 |
| 12 | KC156115 |
| 13 | AY162223 |
| 14 | EU293445 |
| 15 | EU293447 |
| 16 | AF411966 |

**Table 4:** List of the 78 Hepatitis B Virus (HBV) samples chosen from the HBVdb database.

| Sl no | GenBank Accession ID | Sl no | GenBank Accession ID |
| --- | --- | --- | --- |
| 1 | EU859930 | 40 | KP659254 |
| 2 | KU605535 | 41 | JF754609 |
| 3 | KU605536 | 42 | MH464843 |
| 4 | AF143298 | 43 | JN664934 |
| 5 | AB014370 | 44 | AB033558 |
| 6 | AB116077 | 45 | AB104712 |
| 7 | AB116079 | 46 | AB104710 |
| 8 | AB246335 | 47 | AB555501 |
| 9 | DQ020002 | 48 | AB583680 |
| 10 | DQ315785 | 49 | EU594415 |
| 11 | EF103278 | 50 | EU594421 |
| 12 | EF208113 | 51 | FJ349216 |
| 13 | EU054331 | 52 | GQ183473 |
| 14 | EU594389 | 53 | GQ477452 |
| 15 | EU410082 | 54 | GU456657 |
| 16 | EU859948 | 55 | HM750151 |
| 17 | FN545840 | 56 | HQ833466 |
| 18 | FN545836 | 57 | JN040754 |
| 19 | KT749840 | 58 | JN664936 |
| 20 | MK512473 | 59 | KC875283 |
| 21 | AB010289 | 60 | KF471658 |
| 22 | AB033554 | 61 | AB375170 |
| 23 | AB033555 | 62 | HE981171 |
| 24 | AB100695 | 63 | KF414679 |
| 25 | AB205122 | 64 | MT426120 |
| 26 | AB205121 | 65 | AF405706 |
| 27 | AB241117 | 66 | KF767450 |
| 28 | AB246342 | 67 | AB375165 |
| 29 | AB900111 | 68 | AB375166 |
| 30 | AB981581 | 69 | KX264500 |
| 31 | AF282917 | 70 | KM998716 |
| 32 | AP011085 | 71 | KR230749 |
| 33 | AY167100 | 72 | GU563556 |
| 34 | DQ361535 | 73 | KY004111 |
| 35 | DQ975271 | 74 | AB064310 |
| 36 | EU439018 | 75 | EF464098 |
| 37 | EU939673 | 76 | EF464097 |
| 38 | FJ349236 | 77 | EF634480 |
| 39 | GQ377537 | 78 | GU565217 |

**Table 5:** List of the 30 SARS-CoV-2 (B.1.617.1 strain) samples chosen from the NCBI database.

| Sl no | GenBank Accession ID | Sl no | GenBank Accession ID |
| --- | --- | --- | --- |
| 1 | OM915404 | 16 | MZ359879 |
| 2 | OM916775 | 17 | MZ3559881 |
| 3 | MZ724466 | 18 | MZ340525 |
| 4 | MZ562746 | 19 | MZ30541 |
| 5 | MZ562747 | 20 | MZ300543 |
| 6 | MZ562748 | 21 | MZ342807 |
| 7 | MZ562756 | 22 | MZ342889 |
| 8 | MZ562758 | 23 | MZ332524 |
| 9 | MZ562759 | 24 | MZ332527 |
| 10 | MZ558094 | 25 | MZ314697 |
| 11 | MZ021506 | 26 | MZ314701 |
| 12 | MZ401492 | 27 | MZ314702 |
| 13 | MZ401493 | 28 | MZ310587 |
| 14 | MZ401500 | 29 | MZ310590 |
| 15 | MZ363839 | 30 | MZ269263 |

